# Sentence superiority in the reading brain

**DOI:** 10.1101/2023.04.17.534292

**Authors:** Stéphane Dufau, Jeremy Yeaton, Jean-Michel Badier, Sophie Chen, Phillip J. Holcomb, Jonathan Grainger

## Abstract

When a sequence of written words is briefly presented and participants are asked to identify just one word at a post-cued location, then word identification accuracy is higher when the word is presented in a grammatically correct sequence compared with an ungrammatical sequence. This sentence superiority effect has been reported in several behavioral studies and two EEG investigations. Taken together, the results of these studies support the hypothesis that the sentence superiority effect is primarily driven by rapid access to a sentence-level representation via partial word identification processes that operate in parallel over several words. Here we used MEG to examine the neural structures involved in this early stage of written sentence processing, and to further specify the timing of the different processes involved. Source activities over time showed grammatical vs. ungrammatical differences first in the left inferior frontal gyrus (IFG: 325-400 ms), then the left anterior temporal lobe (ATL: 475-525 ms), and finally in both left IFG and left posterior superior temporal gyrus (pSTG: 550-600 ms). We interpret the early IFG activity as reflecting the rapid bottom-up activation of sentence-level representations, including syntax, enabled by partly parallel word processing. Subsequent activity in ATL and pSTG is thought to reflect the constraints imposed by such sentence-level representations on on-going word-based semantic activation (ATL), and the subsequent development of a more detailed sentence-level representation (pSTG). These results provide further support for a cascaded interactive-activation account of sentence reading.

## 1. Introduction

The sentence superiority effect reflects the literate brain’s ability to better recall a grammatically correct sequence of words compared with an ungrammatical sequence. Since its discovery by Cattell (1886, in Scheerer, 1981), it has been used to study the relational binding of words according to certain syntactic and grammatical rules (see Roverud et al., 2020, for an overview). Typically, in such studies, words are presented one after another (serially) either in a visual or auditory modality. Recently Snell and Grainger (2017) asked whether or not such an effect could be observed using a brief visual parallel—rather than serial—presentation of a sentence. In this paradigm—the Rapid Parallel Visual Presentation (RPVP) procedure—a word would be better recognized when it is embedded in a syntactically correct (grammatical) sequence. Such a finding would echo the word superiority effect, whereby a letter is better identified when presented embedded in a word rather than a random string of letters (Cattell, 1886, Reicher, 1969, Wheeler 1970). Snell and Grainger asked their participants to identify a single target word that was embedded either in a 4-word sentence or in an ungrammatical scrambled sequence of the same words. The authors were interested in knowing whether a 200-millisecond display of a word sequence was sufficient for a coarse syntactic structure to emerge, which could influence word identification accuracy via feedback to on-going word identification processes. The results showed that words-in-sentences were recognized with better accuracy (+20%) than words in random sequences, regardless of their position in the sequence. This effect, initially studied in literate adults, was also found in 9-year-old primary school children who showed a smaller but significant accuracy gain of around 10% (Massol & Grainger, 2020).

The sentence superiority effect observed with the RPVP paradigm raises the question as to the mechanisms driving this effect. Snell and Grainger interpreted their finding in an Interactive-Activation framework (McClelland & Rumelhart, 1981) applied to sentence reading, whereby a fast bottom-up activation of sentence-level structures (when these are available) would reinforce word identification processes via top-down feedback from sentence-level structures to words. However, an alternative explanation for the sentence superiority effect obtained with the RPVP paradigm could appeal to the sophisticated-guessing explanation of the word superiority effect (Johnston, 1978). That is, identification of part of the sentence context (one or two words and their association) would enable participants to make informed guesses about the identity of the post-cued target word given partial information about that word (e.g., one or two letters). To put these two interpretations to test, an electroencephalography (EEG) experiment was conducted by Wen et al. (2019) using the same materials and procedure as Snell and Grainger (2017). The authors found a difference in the event-related potential (ERP) traces that started at around 270 ms with the bulk of the sentence superiority effect occurring in an early onsetting N400 component. Such an effect, emerging within the classical time-window of lexical access as revealed by EEG studies (see Grainger & Holcomb, 2009, for a review), pointed to online, automatic processing of linguistic information (word identities and sentence-level structures) rather than offline guessing procedures as the source of the sentence superiority effect (see Wen et al., 2020, for a similar timing of ERP effects in the RPVP paradigm but with a grammatical decision task).

Here, we conducted a similar experiment to Wen et al. (2019) but using magnetoencephalography (MEG) rather than EEG in order to associate the timing of the visual sentence superiority effect with the underlying neural structures. MEG affords much higher spatial resolution than EEG and is thus the gold standard technique for cortical source reconstruction combined with millisecond timing. For the present experiment, we used the same materials and procedure as the two preceding studies (Snell & Grainger, 2017; Wen et al., 2019), however our participants had to report the post-cued target word by reading it out loud rather than typing it using a computer keyboard.

Current findings suggest that the sentence superiority effect is primarily, but not exclusively, driven by syntactic processing (Declerck et al., 2020; Massol et al., 2021). In line with our theoretical work on sentence reading (Snell et al., 2017; 2018), the first phase of processing driving this effect would involve the rapid association of word identities (partially performed in parallel) with the corresponding parts-of-speech, hence enabling the construction of a primitive (or “good-enough”: Ferreira & Lowder, 2016) syntactic structure. The processing of semantic information associated with words would provide further input to the construction of a sentence-level representation (see e.g., Massol et al., 2021) that provides feedback to on-going word identification processes. The overall processing driving the sentence superiority effect is therefore thought to involve partially parallel word identification, the association of word identities with parts-of-speech, the activation of semantic representations associated with word identities, and the construction of a sentence-level representation on the basis of this syntactic and semantic information. Crucially, this processing is thought to be cascaded and interactive in nature, hence allowing a sentence-level representation to be generated rapidly enough to modulate the ease with which a single word in the sentence can be identified.

It is important to note that much prior research investigating the neural structures associated with syntactic processing has focused on spoken language (see Friederici & Kotz, 2003; Matchin & Hickok, 2020, for reviews), whereas the present study investigates written sentence processing with simultaneous presentation of words. This parallel word presentation contrasts with the serial presentation typically used with sequences of written words in EEG studies (e.g., Hagoort & Brown, 2000; Osterhout & Holcomb, 1992), MEG studies (e.g., Halgren et al., 2002; Marinkovic et al., 2003; Matchin et al., 2019), and fMRI studies (e.g., Wilson et al., 2018; Udden et al., 2022; see Walenski et al., 2019, for a review of written sentence processing). Therefore, the regions of interest (ROIs) for data analysis were primarily determined by prior research using written materials presented simultaneously and not serially. Nevertheless, it should be noted that the ROIs selected for the present study do form part of the more general “language network”, highlighted by extensive fMRI research (e.g., Fedorenko et al., 2011; Hagoort, 2014; Price, 2012) and confirming the picture already painted by lesion studies in the 19^th^ century (Broca, 1861; Wernicke, 1874). This language network encompasses neural structures in the left frontal and temporal cortices that include Broca’s area and Wernicke’s area.

In sum, the key contribution of the present work consisted in investigating the processing of written word sequences presented simultaneously (as opposed to the serial presentation typically used in prior EEG, MEG, and fMRI investigations of written sentence processing) and under data-limited presentation conditions (i.e., brief stimulus exposures). We expected these specific testing conditions to help isolate the neural structures involved in the rapid elaboration of a primitive sentence-level representation during written sentence comprehension. On the basis of prior fMRI work with simultaneously presented sequences of written words (e.g., Bonhage et al., 2015; Pallier et al., 2011; Schuster et al., 2016) we expected to find activation related to sentence-level processing predominantly in the left inferior frontal gyrus (IFG) and left posterior temporal regions (pSTG, pMTG), as well as activity in the anterior temporal lobe (ATL: Carter et al., 2019; Henderson et al., 2016). On the basis of our prior ERP work (Wen et al., 2019), we expected to observe differences between sentences and ungrammatical sequences to first emerge around 300 ms post-stimulus onset (i.e., onset of the complete sequence of words), and to cover the epochs traditionally associated with the N400 and P600 ERP components. Given the hypothesized early role for syntax in our proposed account of the sentence superiority effect, we expected to observe the earliest differences in left IFG given the evidence that IFG plays a key role in processing syntactic information during spoken language comprehension (e.g., Hagoort, 2014; Friederici, 2017; Friederici & Kotz, 2003) and text comprehension under conditions of parallel presentation (Constable et al., 2004; Henderson et al., 2016). This was then expected to be followed by activity in the left anterior temporal lobe (ATL) where it is thought that syntactico-semantic integration occurs (Carter et al., 2016), and where sentence-level representations begin to constrain on-going word identification (Carter et al., 2019; Henderson et al., 2016). We then expected to observe differences in left posterior temporal regions (pSTG, pMTG), regions that are traditionally associated with the elaboration of more detailed syntactic representations for speech and text (Constable et al., 2004). Finally, we expected to observe later differences in pSTG, driven by processes that combine lexico-semantic and syntactic information for sentence comprehension (e.g., Friederici & Kotz, 2003), as thought to be reflected in the P600 ERP component (Osterhout & Holcomb, 1992).

## 2. Materials and methods

Material and methods exposed below closely follow those of the reported studies conducted in the MEG center of La Timone Hospital, Marseille, France.

### 2.1. Participants

Twenty native French speaking adults (mean age= 23.65 years; range= [20;30]; SD= 3.01; 14 women; 6 men) were included in this research and earned EUR 50 for participation. All the participants were recruited from Aix-Marseille University (France). In order to prevent adding unnecessary noise in the MEG signal, participants with non-removable piercings or surgical implants were not included in this study. Participants were asked not to wear metallic or magnetic jewelry and to avoid using cosmetic products. Participants were monolingual French native speakers, had no record of neurological or psychiatric disorders and had regular or corrected hearing and vision. This study was carried out in accordance with the recommendations of the French ethics committee (Comité de Protection des Personnes SUD-EST IV No. 17/051). All participants gave written informed consent in accordance with the Declaration of Helsinki.

### 2.2. Stimuli and Design

Materials from Snell and Grainger (2017) were used in this experiment. They consisted of four-hundred 4-word sequences that were either syntactically correct or not (word length, average= 4.03, SD= 0.82 letters; word frequency, average= 5.66, SD= 1.08 Zipf units^1^; 200 syntactically correct sequences). The set of sequences was built in the following way. First, two hundred syntactically correct sequences were constructed and tested using cloze probability measures to ensure that they were semantically neutral (for details, see Snell & Grainger, 2017). Syntactically incorrect sequences were then derived from the correct pool: each correct sequence was associated with an incorrect sequence that was composed of the same words but in a different order. Within each pair of sequences, a word was chosen as the target (used in the identification task; counter-balanced across trials using the 4 possible word positions) and remained in the same position within the two sequences while the three other words were randomly permuted to form the incorrect sequence.

This procedure ensured that the obtained scrambled sequence was not syntactically correct while keeping intact the properties of the individual words composing the pair of sequences. To avoid within-participant stimulus repetition, two counterbalanced lists of 200 sequences each (100 syntactically correct and 100 syntactically incorrect sequences) were created, and participants were assigned to one of them. As such, all stimuli were presented in both syntactic conditions (across two participants) and each participant was presented a single member of a sequence pair. Each list of 200-word sequences was divided into 4 blocks of 50 sequences. A 2 (Grammatical vs. Ungrammatical) x 4 (within-sequence word target position) factorial design was used, and stimulus order was randomized within blocks.

### 2.3. Procedure

We used E-Prime software (Psychology Software Tools, 1999) to implement a visual post-cued word-in-sequence identification task where 4 horizontally aligned words were briefly presented (RPVP paradigm). Each trial began with the visual presentation of a fixation sign in the centre of the screen consisting of 2 bars vertically separated by an empty space (see Figure 1). After 500 milliseconds (ms), the empty space was filled by a sequence (either syntactically correct or incorrect) for 200 ms. The sequence’s letters were then replaced by a post-mask formed of hash symbols (#) preserving between-word spaces and inter-letter spacing. The post-mask consisted of two colors: the hashes replacing the target word were displayed in yellow and the three other locations were displayed in white. The task consisted of naming the word which had appeared in the cued (yellow) position. The stimulus remained on screen until a verbal response was given. When E-Prime detected an oral response, the visual sequence disappeared, leaving an empty screen for 100 ms followed by a stimulus composed of a single letter “C” (from the French word *cligner* – to blink) in the middle of the screen presented for 1500 ms. A trial ended with an empty screen for 100 ms. Participants were instructed to fixate the empty space in between bars at the beginning of the trial and to blink only during the “C” phase of a trial. Vertical bars remained on screen from the start of the trial to the display of the letter C. All visual stimuli were displayed using a white 18pt bold Courier New font on a black background.

**Figure 1.**
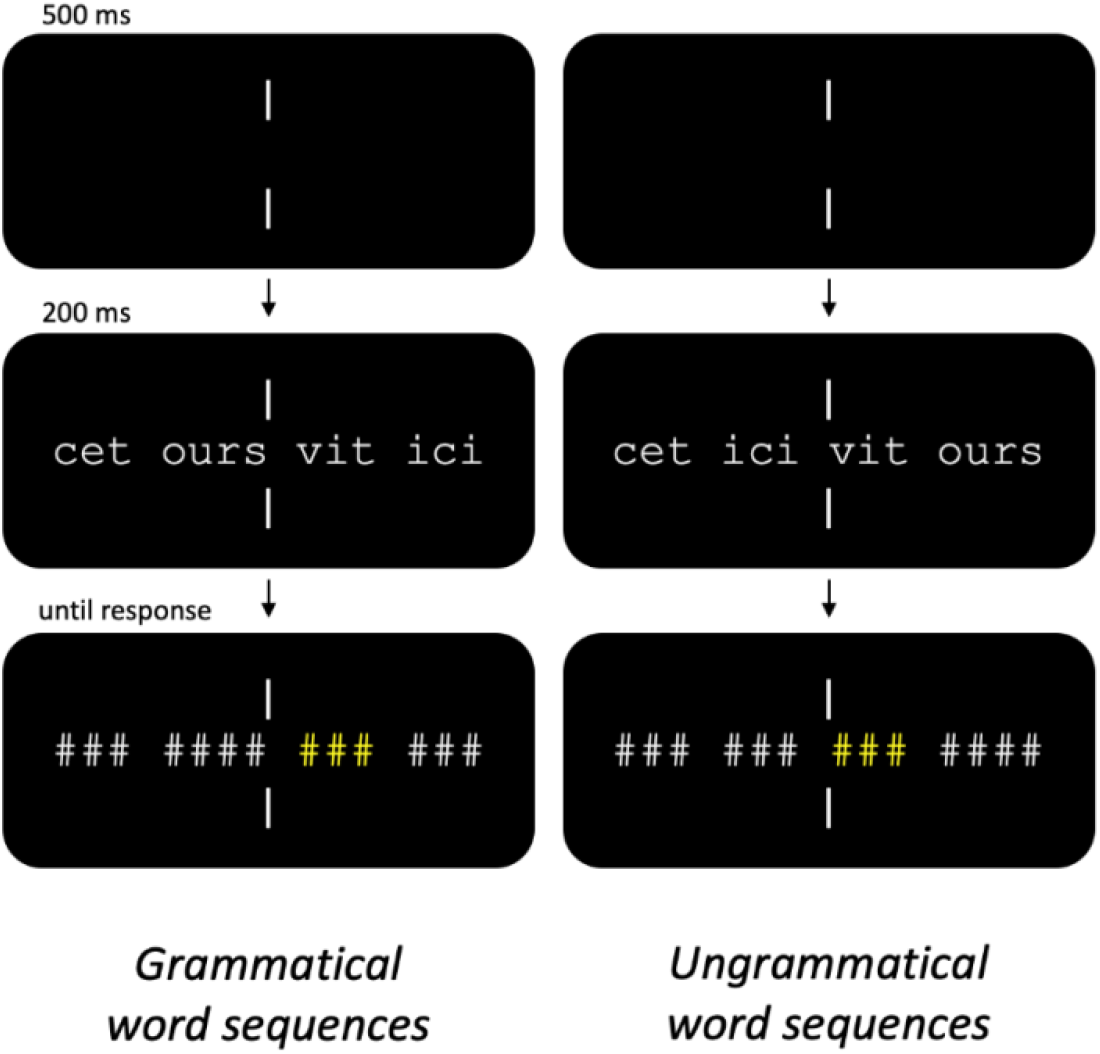
Trial description for the grammatical (left panel) and ungrammatical (right panel) word sequences. After an initial fixation stimulus was displayed for 500 milliseconds (ms; see Methods section 2.3), a word sequence appeared that was either syntactically correct (*cet ours vit ici* - this bear lives here) or not (*cet ici vit ours* - this here lives bear). The word sequence was visible for 200 ms.

### 2.4. Data acquisition and analysis

The data were acquired on a 248-channel whole-head 4D Neuroimaging MEG system at La Timone Hospital in Marseille, France (4D Neuro-imaging, San Diego, CA). Sampling rate was 2035 Hz. Individual head shapes, consisting of the forehead, nose, and the location of the head-position coils were digitized using a 3D Polhemus Fastrak device (Polhemus Inc., Colchester, VT, USA). Five head-positioning coils were attached to the forehead and periauricular points to determine the position of the head. Head position was captured at the beginning of the first and third block to check and possibly compensate for differences in head position between the first and second half of the stimuli set. Participants were lying inside a magnetically shielded room and were instructed to move as little as possible. Stimuli were presented on an 1024×768 resolution video-projector screen placed about 40 cm in front of participants. The exact timing of stimuli onset was captured using a photodiode that detected brightness changes on the presentation screen. A non-magnetic microphone (Sennheiser MO2000) was placed nearby the participant’s mouth (without skin contact) to capture verbal responses. A stereo audio file was recorded during the entire experiment, with one channel dedicated to the participant’s verbal output and another one to record clicks corresponding to the word sequence display onset timing. Verbal output was coded off-line for the correctness of the response. Following Gross et al. (2013), an electrooculogram was recorded throughout the experiment to capture the activity of eye movements as well as an electrocardiogram for heartbeats.

For all participants but three, cortical surface extraction was performed on individual high-resolution 3D T1-weighted MRI structural images using BrainSuite (Shattuck & Leahy, 2002). For the three participants having a non-conclusive extraction, sources were constrained to a cortical surface mesh template obtained from the MNI ICBM152 brain. Brainstorm was used with default parameters to warp the template to each participant’s digitized head shape. Cortical surface was defined with 15,000 vertices and the MRI was realigned on three fiducials (nasion, left and right pre-auricular points).

Continuous data processing was performed using Anywave (Colombet et al., 2015) for visual rejection of channels showing excessive noise, muscle, or SQUID jump artifacts, for filtering (1-to 300-Hz bandpass) and independent component analysis (runica algorithm) to identify and remove the heartbeat and blink artifacts. Brainstorm (Tadel et al., 2011) was used for additional filtering (25-Hz lowpass), for epoching signal segments time-locked to word sequence onset (−200 to 800 ms), trial rejection and averaging. Data processing and source reconstruction were performed independently for each participant. Artifact-free epochs were averaged separately for each experimental condition to obtain event-relate fields (ERFs) for each participant. This average was then projected onto the cortical surface using a free orientation, cortically constrained minimum norm estimation (MNE) procedure (Hauk, 2004; Hämäläinen & Ilmoniemi, 1994). The MNE was weighted by a sample estimate of sensor noise covariance matrix obtained from empty room recordings for each of the participants and used for improved data modelling as is typical in MNE approaches (Baillet et al., 2001; Dale et al., 2000). The MEG forward model was obtained from overlapping spheres fitted to each participant’s scalp points (Huang et al., 1999). For all participants but three, cortical surface extraction was performed on individual MRI images using BrainSuite (Shattuck & Leahy, 2002). For the three participants having a non-conclusive extraction, sources were constrained to a cortical surface mesh template obtained from the MNI ICBM152 brain. Brainstorm was used with default parameters to warp the template to each participant’s digitized head shape. The norm of the three source time series at each cortical voxel (i.e., conversion of orientation-unconstrained sources to flat maps, taking the norm of the three elementary dipoles at each time step, yielding only one value by vertex) was extracted. The difference was taken between the source projections for the two conditions for each participant, then the difference was z-scored with respect to the [−200, 0] milliseconds baseline, and absolute value transformed, thus yielding a single value corresponding to the difference in activation between conditions by vertex at each time point.

We defined four regions of interest (ROIs; see Figure 2) corresponding to the left hemisphere brain areas discussed in the Introduction: inferior frontal gyrus (IFG; x=47, y=143, z=85, N=287 vertices), posterior superior temporal gyrus (pSTG; x=30, y=90, z=93, N=227 vertices), anterior temporal lobe (ATL; x=28, y=121, z=58, N=184 vertices), and posterior middle temporal gyrus (pMTG; x=29, y=92, z=70, N=144 vertices). To obtain the time course of activity in each ROI, the average difference between the conditions (described above) was taken over all the vertices comprising that region. This ROI time course was then compared to the baseline using a t-test procedure^1^ (one-tail, positive) with an alpha level of 0.05, and a Bonferroni correction for multiple comparisons over the time dimension. Only regions with at least 50ms continuously above the alpha threshold were considered significant.

**Figure 2.**
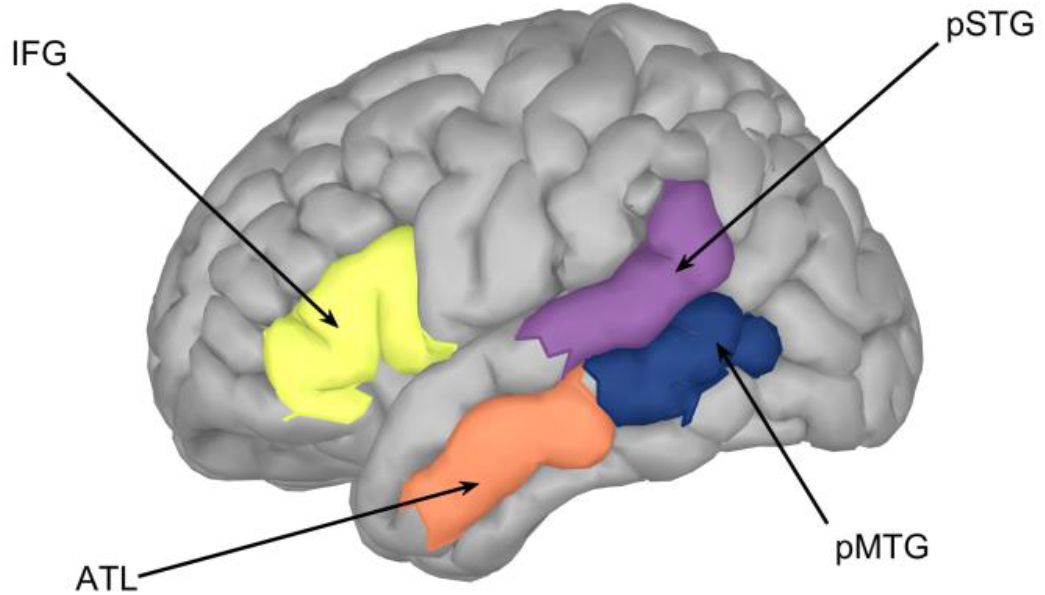
Illustration of the ROIs selected for their noted role in sentence-level processing in the left hemisphere of the human brain. IFG: inferior frontal gyrus; pSTG: posterior superior temporal gyrus; ATL: anterior temporal lobe; pMTG: posterior middle temporal gyrus.

## 3. Results

Accuracy results of the word-in-sequence identification task are shown in Figure 3. The accuracy data were analyzed in a 2 (Grammatical, Ungrammatical) x 4 (Positions) logistic mixed-effects regression modeling (Jaeger, 2008). Overall, individual words were better identified when the words were presented in a syntactically-correct sequence (62.4% for Grammatical, 50.3% for Ungrammatical), b=0.83, SD=0.36645, z=2.26 (Pr(>|z|) = 0.024. The effect of Position and the interaction between Position and Grammaticality were not significant.

**Figure 3.**
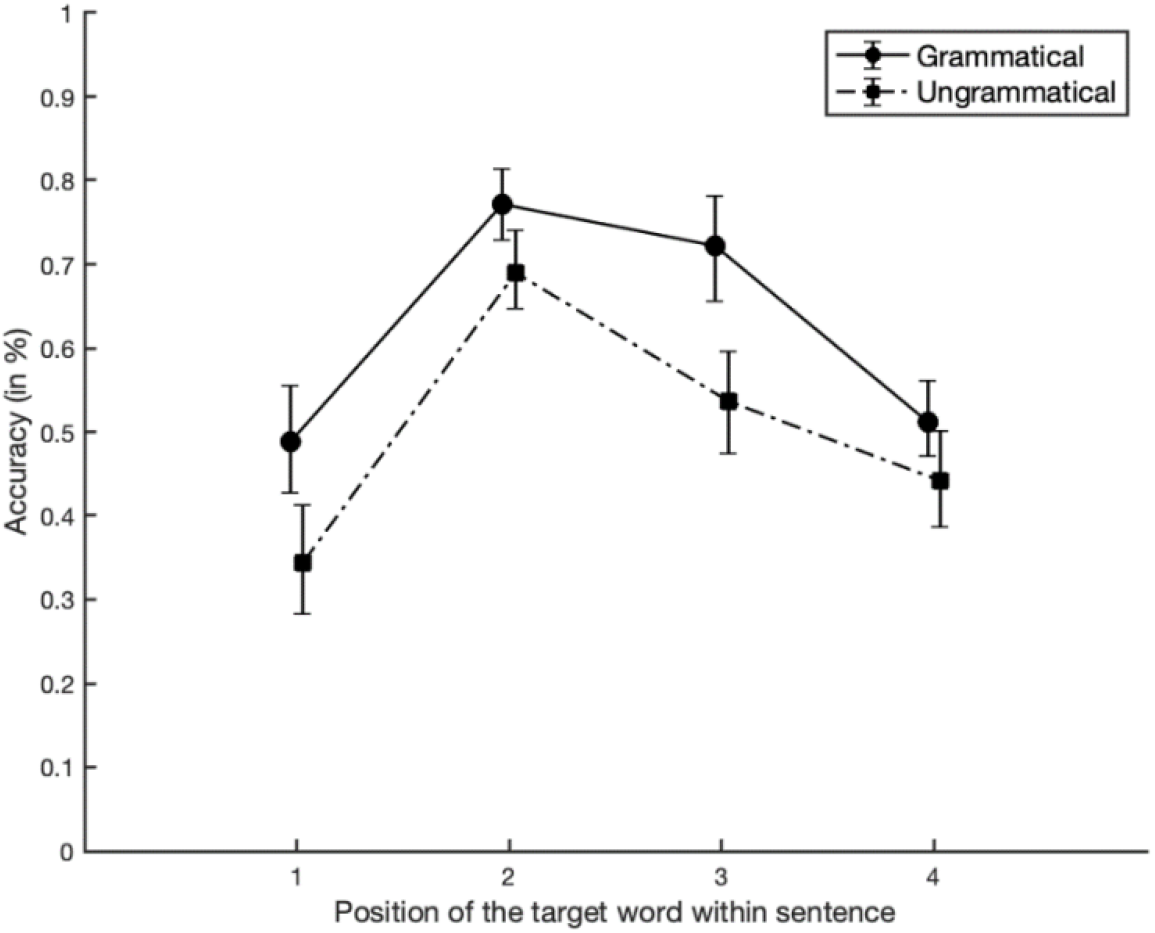
Mean accuracy per position of the target word (1: most leftward word of the horizontal sequence; 4: most rightward). Responses in the Grammatical condition (words in grammatical sequences) had a higher accuracy than responses in the Ungrammatical condition (words in ungrammatical sequences) at all positions. Error bars correspond to the bootstrapped by-participant 95% confidence interval standardized for participants across positions (Cousineau, 2005).

Cortical source activities for the Grammatical and Ungrammatical conditions in 4 ROIs in left hemisphere (inferior frontal gyrus, IFG; posterior superior temporal gyrus, pSTG; anterior temporal lobe, ATL; posterior middle temporal gyrus, pMTG) were compared using a Bonferroni-corrected t-test at each time point. Three out of the four ROIs revealed a significant difference (see Figure 4): IFG in 2 time-windows (321-406 ms; 549-602 ms), ATL in a single time window (466-531 ms), and pSTG in a single time window as well (553-622 ms).

**Figure 4.**
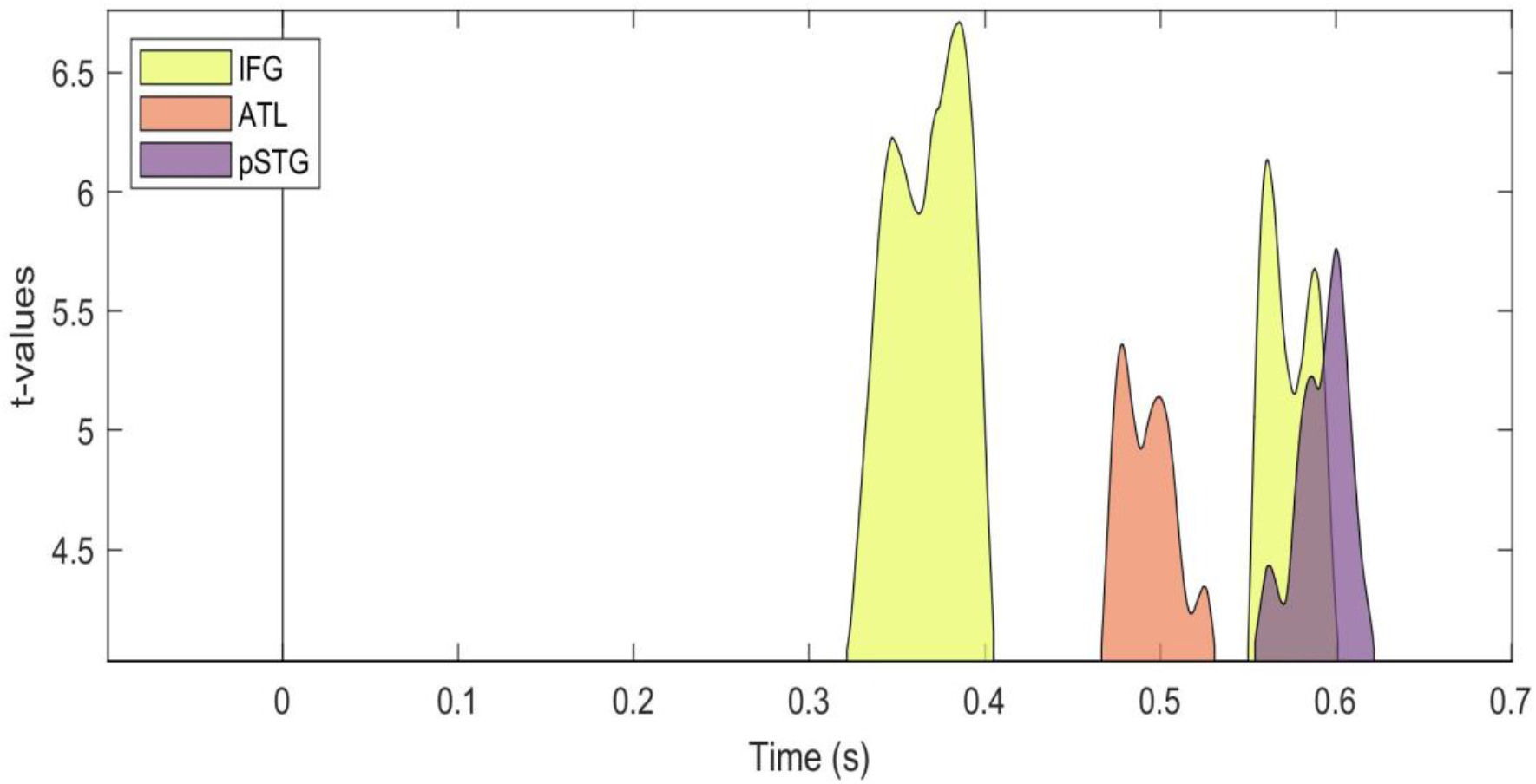
Results of a t-test comparing the Grammatical-Ungrammatical difference against baseline for the 4 ROIs in the [-100; 700] ms time window across participants (Bonferroni-corrected; minimum of 50 milliseconds of continuous positivity). Y-axis scale for t-values was cut at significance value (∼4.1).

## 4. Discussion

Presented in the context of a sequence of written words, a word is better recognized when the sequence is grammatical (i.e., within a sentence or a phrase) rather than an ungrammatical word sequence. This sentence superiority effect was recently brought to light in a series of behavioral and EEG experiments that used post-cued partial report with the Rapid Parallel Visual Presentation (RPVP) procedure (e.g., Snell & Grainger, 2017; Wen et al., 2019). In these studies, enhanced word identification was observed in the grammatical compared with the ungrammatical sequences. Here, we used the materials and procedure of Snell and Grainger (2017) in an MEG setting to study the spatio-temporal dynamics of the sentence superiority effect. As expected, behavioral results revealed the standard sentence superiority effect (i.e., word identification accuracy was greater in the grammatical sequences compared with the ungrammatical sequences). This pattern is evidence that some form of primitive sentence-level representation had been computed even with the very brief (200 ms) stimulus exposures used in the RPVP paradigm. Based on prior fMRI research investigating the processing of simultaneously presented sequences of written words (Bonhage et al., 2015; Carter et al., 2019; Henderson et al., 2016; Pallier et al., 2011; Schuster et al., 2016) we identified 4 ROIs as brain regions likely to be involved in written sentence processing, and we examined the sentence superiority effect in each of these ROIs (all located in the left hemisphere). Source activations over time revealed grammatical vs. ungrammatical differences in three of these ROIs: first in IFG, then ATL, and finally in both IFG and pSTG (see Figure 4). We now examine in more detail the potential role of these three ROIs in subserving the sentence superiority effect, and we discuss the implications of the relative timing of the activities that we observed.

### Early IFG activity

A previous EEG investigation using the same materials as in the present study found an early onsetting effect in the N400 time-window (274-410 ms) where ungrammatical sequences produced more negative-going waveforms than the grammatical condition (Wen et al., 2019). This timing aligns with the first significant activity we found in the IFG (321-406 ms), and therefore, following our interpretation of the EEG findings, points to a key role for IFG in rapidly generating a primitive sentence-level representation upon presentation of a sequence of written words. We suspect that this primitive sentence-level representation takes the form of a “good-enough” syntactic structure, as hypothesized by Ferreira and Lowder (2016; see Friederici, 2011, for an earlier proposal for spoken language comprehension). Further evidence for an early role of IFG in written sentence processing, albeit obtained with a serial presentation procedure, was reported in the MEG study of Halgren et al., (2002). In that study the comparison was between sentences ending in a semantically congruous or incongruous word. In line with our results, activity in IFG was found starting around 300 ms post final word onset.

### ATL activity

We expected to see activity in ATL given prior fixation-related fMRI work suggesting that this could be the locus of the initial phase of integration of syntactic and semantic information for sentence comprehension (Carter et al., 2016) and where sentence-level representations begin to constrain on-going word identification (Carter et al., 2019; Henderson et al., 2016). The timing of the observed activity in ATL (475-525 ms) fits with this interpretation, according to which an early primitive syntactic representation is first computed by IFG (as attested by the present study and the EEG study of Wen et al., 2019) followed by the combination of this syntactic information with on-going semantic processing in ATL as a key step in interpreting the sentence being read. Once again, these findings fit with the earlier MEG study of Halgren et al. (2002; see also Brennan & Pylkkänen, 2012; Matchin et al., 2019) that revealed activity in ATL in the time-window of the N400 ERP component, hence pointing to a role for semantics in generating activity in ATL.

### pSTG activity co-occurring with a second phase of activity in IFG

The final structures that were revealed by our sentence superiority manipulation (pSTG and IFG, between 550-600ms) point to an interactive process involving the primitive syntactic structure generated by IFG and the construction of a more detailed sentence-level representation that combines both syntactic and semantic information, but going beyond what is already achieved by ATL. As already proposed by Friederici (2017), on the basis of spoken language studies, pSTG could well be the host of syntactic reanalysis and repair. In other words, activity in IFG followed by ATL would enable the generation of a tentative sentence-level representation combining syntax and semantics, and the later activity in both pSTG and IFG would reflect a process of verifying that the sentence-level structures computed by IFG and pSTG are compatible. The relatively late activity found in STG in the Halgren et al. (2002) MEG study could also reflect such a verification process, which would be triggered by both syntactic and semantic anomalies. Very tentatively, this might well involve a top-down matching of the more detailed sentence-level representation (syntax and semantics) with the word identities and parts-of-speech that are thought to form the basis of a primitive sentence-level structure.

## 5. Conclusions

In an MEG investigation of the sentence superiority effect, we found source activities revealing significant differences between the grammatical and ungrammatical word sequences first in the left IFG, then the left ATL, and finally in both left IFG and left pSTG. We interpret the early IFG activity as reflecting the rapid bottom-up activation of sentence-level representations from a sequence of written words, and in particular a primitive syntactic representation of the sequence of words when that is available. Subsequent activity in ATL and pSTG is thought to reflect the constraints imposed by the early sentence-level representation on on-going word-based semantic activation and the integration of semantic and syntactic sentence-level information (ATL), followed by the subsequent development of a more detailed sentence-level representation (pSTG), including verification that this more detailed representation is compatible with the early primitive representation computed by IFG.

## FOOTNOTES

1. Comparison was made on the mean over all vertices in the ROI.

## Data/code availability statement

Stimuli and data are available at https://osf.io/wzq5y/.

## Ethics statement

Ethics approval was given by the French ethics committee (Comité de Protection des Personnes SUD-EST IV) agreement No. 17/051. All participants gave written informed consent in accordance with the Declaration of Helsinki.

## Author contributions

Conceptualization (JG); Data curation (SD, JY, JMB, SC); Formal analysis (SD, JY); Funding acquisition (JG); Investigation (SD, SC); Methodology (SD, JMB, SC, JG); Project administration (SD, JMB); Resources (JMB, SC); Software (SD, JY); Supervision (SD, JMB, SC, JG); Validation (SD, JY, JMB, SC, PH, JG); Visualization (JY); Roles/Writing - original draft (SD, JY, JG); Writing - review & editing (SD, JY, PH, JG).

